# Loss of tumor suppressor p53 upregulates stem cell factor SOX9 via Notch signaling

**DOI:** 10.1101/2025.05.27.656408

**Authors:** Rahul Sanawar, Wenjun Guo

**Affiliations:** Department of Cell Biology, Albert Einstein College of Medicine, Bronx, NY 10461, USA; Ruth L. and David S. Gottesman Institute for Stem Cell and Regenerative Medicine Research, Albert Einstein College of Medicine, Bronx, NY 10461, USA; Cancer Dormancy Institute, Albert Einstein College of Medicine, Bronx, NY 10461, USA; Marilyn and Stanley M. Katz Institute for Immunotherapy for Cancer and Inflammatory Disorders, Albert Einstein College of Medicine, Bronx, NY 10461, USA; Montefiore Einstein Comprehensive Cancer Center, Albert Einstein College of Medicine, Bronx, NY 10461, USA

## Abstract

Basal-like breast cancer (BLBC) consists of the majority of triple-negative breast cancer subtype that has a higher degree of cellular plasticity owing to a greater number of stem-like cancer cells compared to other subtypes ^1^. BLBCs are thought to originate from the luminal progenitor cells despite their prominent basal-cell features. SOX9 is a key transcription factor which is expressed selectively in estrogen receptor-negative luminal progenitors in postnatal mammary glands. During BLBC progression, SOX9 upregulation is required for the de-differentiation of luminal cells to multipotent fetal mammary stem cell-like states, which are crucial for the malignant progression of BLBC. However, the mechanism driving SOX9 upregulation in BLBC remains unclear. Since p53 is mutated in nearly 90% of the BLBC and is considered as an early event in the BLBC transformation, we hypothesized that p53 loss could contribute to SOX9 upregulation during BLBC progression. Using primary mammary cell organoids, we showed that p53 loss not only induced the SOX9 expression but also stabilized the half-life of SOX9 protein. We further identified that p53 loss increased Psen2 expression to activate the Notch1 signaling, which induced SOX9 expression. Together, our work identified a molecular mechanism allowing loss of tumor suppressor p53 to coopt the cell fate determinant SOX9 to drive de-differentiation in BLBC.

## Introduction

Basal-like breast cancer (BLBC) is considered one of the most aggressive breast cancer subtypes, and these tumors often have high degrees of intratumor cell-state heterogeneity. The underlying cause of such heterogeneity remains poorly understood. Breast tissue is composed of distinct lineages, including two types of luminal cells that are either positive or negative of estrogen receptor (ER) expression and the ER-negative basal cells. While a common fetal mammary stem cell can generate all three lineages, in postnatal mice, each lineage can be maintained by long-lived lineage-restricted stem/progenitor cells, which mediate postnatal mammary gland development and homeostasis ^2-6^. Despite sharing substantial gene expression signatures with basal cells, BLBC is likely to originate from ER-negative luminal cells ^7-12^. Studies showed that the global gene expression signature of BLBC more closely mimics adult luminal progenitors than basal cells, and inactivation of BLBC tumor suppressor BRCA1 in luminal progenitors but not basal cells leads to BLBC formation ^7, 8^. But why luminal progenitors are biased for BLBC development remains an open question.

Mutations in *TP53, RB1, PIK3CA*, and *BRCA1* have been linked with lineage plasticity in multiple cancers ^10, 13-16^. In BLBC, *TP53* is mutated in nearly 90% of the cases ^17-19^. A considerable amount of literature has described the association of *TP53* and/or *RB1* mutations with lineage plasticity in mammary tumors ^20-25^. However, whether and, if so, how these mutations coopt the mammary lineage fate determinants to promote lineage plasticity remain unknown. SOX9, a key developmental transcription factor of the luminal progenitor cells in mouse and human breast ^6, 26^, is required for BLBC initiation from luminal progenitors, as well as for the BLBC progression^27 28, 29^. SOX9 expression has also been linked to cancer progression for various cancer types such as lung adenocarcinoma ^30, 31^, colon carcinoma ^32^, hepatocellular carcinoma ^33^, prostate cancer ^34, 35^, gastric cancer ^36^, non-small cell lung cancer (NSCLC) ^37^, pancreatic cancer ^38-40^, and oral squamous cell carcinoma ^41^, in addition to breast cancer ^26, 42-46^. Using the C3/Tag BLBC mouse model, we have previously shown that SOX9 expression is upregulated in a subset of ER-negative luminal cells during BLBC initiation and that its expression is required for the progression of precancerous mammary intraepithelial neoplasia to malignant BLBC ^27^. Given that C3/Tag transgene induces tumors by inactivating p53, we asked the question whether p53 loss directly induces the expression of lineage plasticity driver SOX9 to promote BLBC transformation.

## RESULTS

### Loss of p53 increases SOX9 expression in mammary organoids

We first analyzed the correlation of p53 status and SOX9 expression in human breast cancer datasets. We found that, in the TCGA dataset, SOX9 expression is significantly higher in patients with p53 mutations (**Fig 1A**). Since p53 is one of the most mutated genes in human breast cancer, we asked the question whether p53 regulates SOX9 expression. Using Sox9-GFP transgenic mice (in which GFP expression is driven by the SOX9 enhancer and promoter) and C3/Tag mice model, we have previously shown that p53 inactivation leads to higher SOX9 expression in the C3/Tag mammary gland than normal control ^27^. However, a functional causality has not been examined.

**Figure 1.**
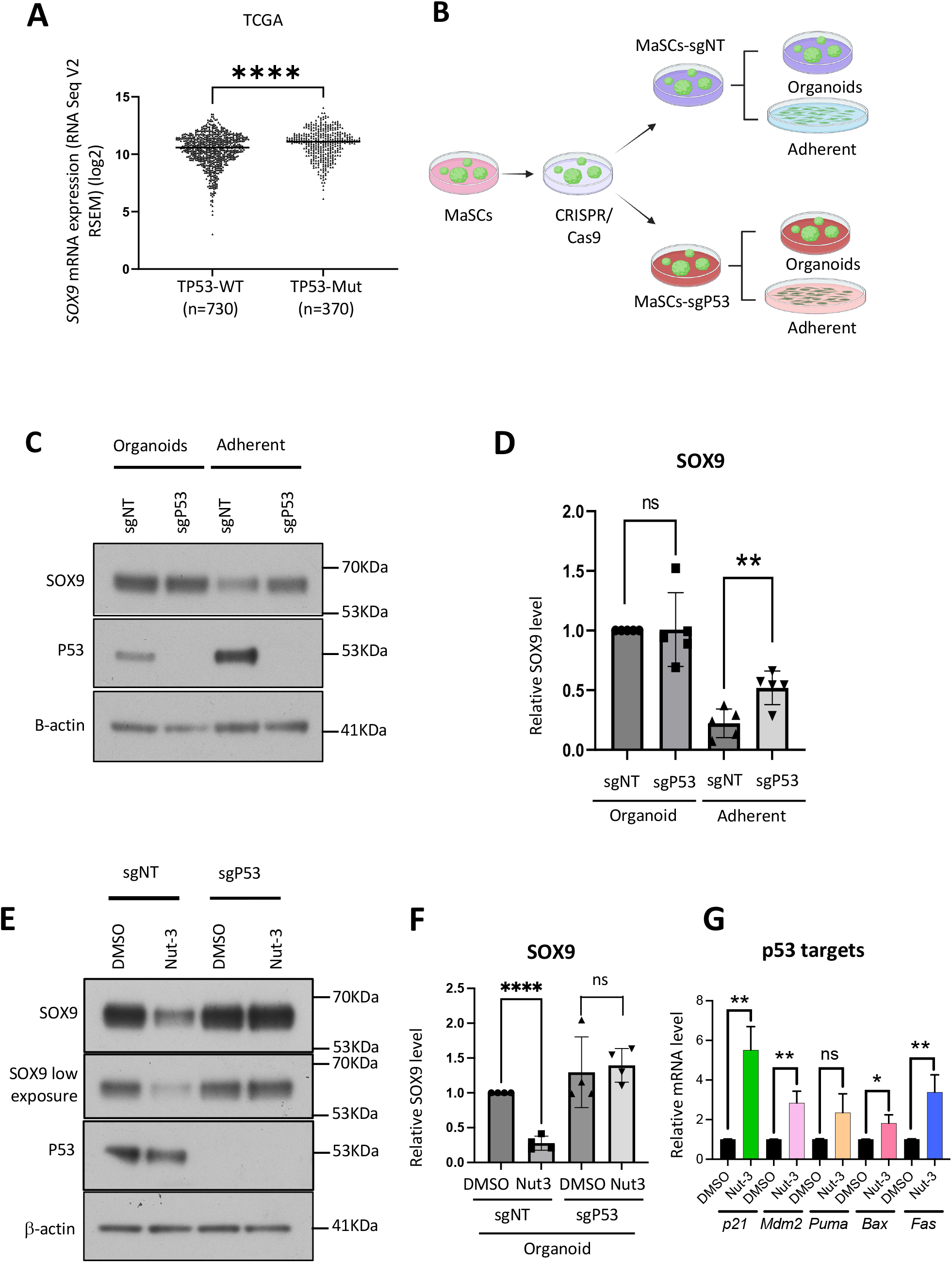
Loss of p53 upregulates SOX9 expression. A. Scatter plots showing the SOX9 mRNA levels among patients stratified into two groups based on the p53 status. Analysis performed on the TCGA dataset (Firehose Legacy, n=982). B. Schematic diagram of the p53 KO experimental flow. The p53-WT and p53-KO MaSCs were grown in adherent condition or organoid condition for 3.5 days before harvesting the cells for RNA or protein isolation. C. WB analysis of p53 and SOX9 in the organoids proficient or deficient for p53. Organoids were cultured in poly-HEMA coated plates in organoid media or in TC coated normal 60mm culture dishes in organoid media without Matrigel for adherent condition (2D). For 3D culture, 300K cells/well of poly-HEMA coated plates were seeded whereas for 2D culture, 150K cells were seeded in 60mm dishes. MaSCs were grown for 3.5 days before isolating proteins and WB. D. Summarized result of the five experiments used for WB in Fig 5C. E. WB analysis measuring the effect of p53 upregulation by Nut-3 on SOX9 expression. p53-WT and p53-KO organoids were treated with DMSO (as control) or 10uM Nut-3 for 48hrs (n=5) in organoid culture condition. 400K MaSCs were admixed with 2ml organoid media with a final conc. of 10uM Nut-3 and 5% Matrigel and seeded in one well of poly-HEMA coated 6-well plates. After 48hrs of treatment, organoids were analyzed by WB. E. Summarized result of four experiments used for WB in Fig 1E. F. RT-qPCR measuring the effect of Nut-3 treatment on p53 downstream targets.

To experimentally check the potential regulatory effect of p53 on SOX9, we first employed p53 loss-of-function approach. We knocked-out (KO) p53 from mammary stem cell (MaSC) organoids ^47^ using lentiviral CRISPR/Cas9 gene targeting (**Fig1B)**. Loss of p53 was validated both by RT-qPCR and western blot (WB) (**Fig1C** and **S1A**). We then assessed the effect of p53 KO on SOX9 in mammary organoids by WB and RT-qPCR (**Fig1C, 1D, S1A and S1B**). We found that p53 KO does not affect the SOX9 mRNA level. Additionally, we did not see any robust change on SOX9 protein levels upon p53 loss, when grown in organoid condition. One possible reason for this could be due to the already very high, potentially saturated, expression of SOX9 in the organoid condition ^48, 49^. Therefore, we examined the effect of p53 loss on SOX9 by switching the mammary organoids to adherent culture condition for 3.5 days. The rationale for this is to downregulate the level of SOX9 substantially by growing the organoid cells in adherent condition so that any regulatory effect on SOX9 protein could be observed more readily. Indeed, mammary cells in adherent culture had significantly lower SOX9 than cells in organoid culture (**Fig 1C**). Interestingly, we found that in adherent condition, p53 loss maintains the SOX9 protein levels closer to what we see in organoids, suggesting loss of p53 induces SOX9 expression (**Fig 1C**). To further confirm this, we performed p53 gain-of-function experiment. For this, we treated the p53-WT and p53-KO organoids with Nutlin-3, an MDM2 inhibitor stabilizing p53. As expected, Nut-3 treatment led to the decreased expression of SOX9 protein in p53-WT organoids (**Fig 1E, 1F**). Nut-3 also significantly upregulated multiple p53 targets (**Fig 1G**). Importantly, Nut-3 did not affect SOX9 levels in p53-KO organoids, indicating Nut-3 acts by stabilizing p53 to downregulate SOX9 expression (**Fig 1E, 1F**). The *SOX9* transcript level was not affected by Nut-3, suggesting the effect of p53 on SOX9 is post-transcriptional (**Fig S1C**).

### Notch induces SOX9 expression

We next investigated which signaling pathway could induce SOX9 expression. Studies have shown various signaling pathways involved in the upregulation of SOX9 expression in different tissue types, including Wnt signaling ^50-65^, FGF10 signaling ^66-73^, TGFβ signaling ^74-76^ and hypoxia signaling ^77-80^. We tried to first access the effects of these signaling pathways on SOX9 in our MaSC organoid system. Surprisingly, we found that activation of these signaling pathways could not increase SOX9 levels in the mammary organoids (**S2A and S2B**).

We then turned our attention to Notch, as SOX9 is reported to be a Notch target in certain tissue types ^81, 82^, and Notch signaling is preferentially activated in the mammary luminal progenitors which express SOX9 within the normal mammary ductal tree ^83, 84^. To test whether Notch signaling regulates SOX9, we treated mammary organoids with DAPT (a γ-secretase inhibitor blocking Notch activation). Treatment with DAPT reduces the Notch signaling activation as assessed by decreased expression of NICD1 (Notch 1 Intracellular Domain) by immunoblot, as well as the decreased expression of the Notch canonical downstream targets, *Hes1* and *Hey1* assessed by RT-qPCR (**Fig 2A** and **2B**). Interestingly, we also observed the decreased expression levels of SOX9 protein as well as transcript in the MaSCs treated with DAPT (**Fig 2A** and **2B**). Next, we cross-validated this observed positive regulatory effect of Notch signaling on SOX9 by activating the Notch signaling in luminal breast cancer cell line, MCF-7. We stably overexpressed NICD1 in MCF-7 cells using a lentiviral vector. NICD1 overexpression leads to a markedly increased expression of SOX9 protein as well as transcript (**Fig 2C** and **2D**). Additionally, we checked this effect in TNBC cell lines, MDA-MB-231, MDA-MB-436 and SUM159. We observed that Notch signaling activation in the TNBC lines also led to an increased expression of SOX9 protein as well as transcript in general (**Fig 2E** and **2F**). Taken together, our findings suggest that Notch signaling induces the expression of SOX9 in normal mammary organoids and breast cancer cells.

**Figure 2.**
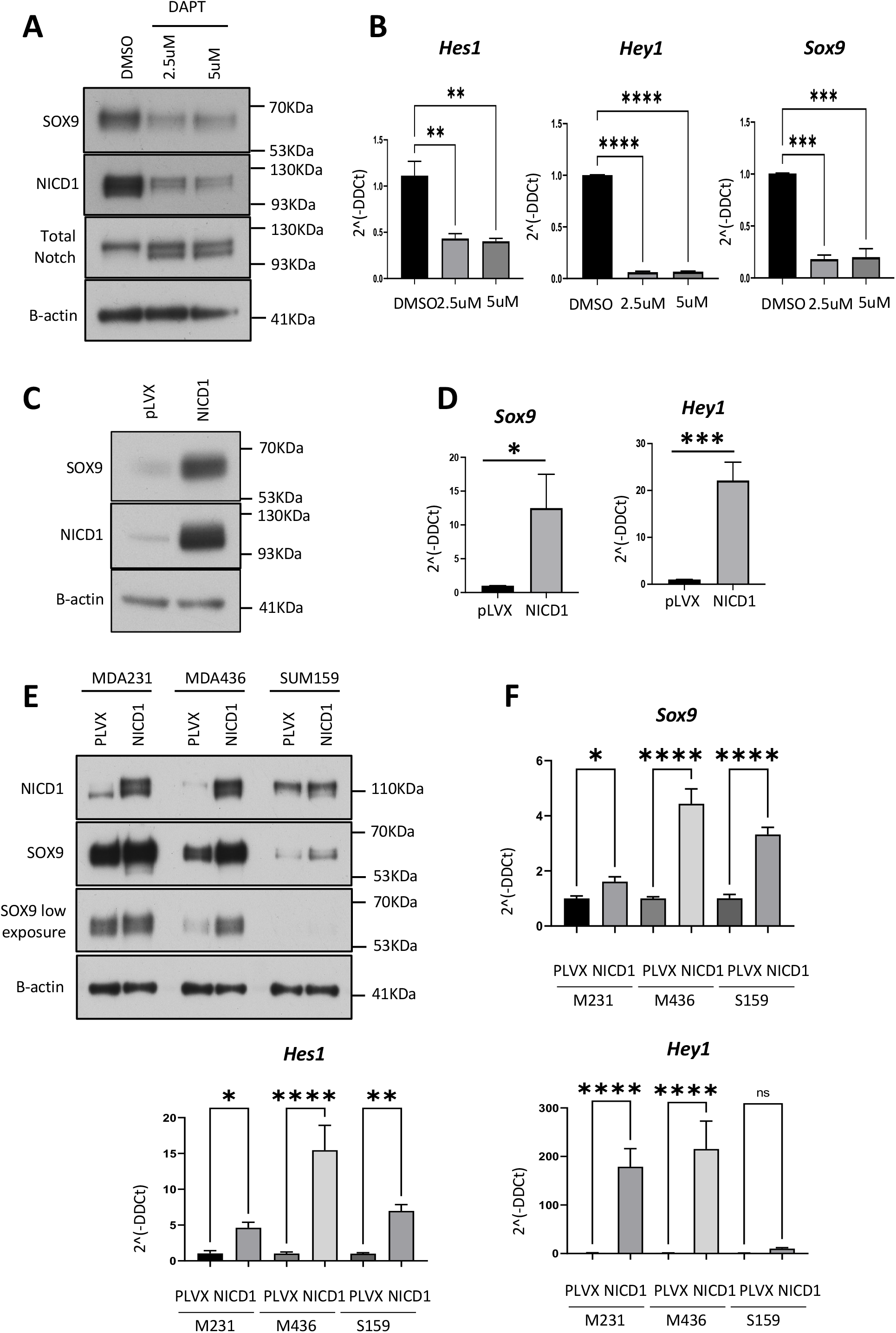
NOTCH signaling upregulates SOX9. A. WB analysis showing the effect of Notch signaling inhibition on SOX9. Organoids were treated with 10uM DAPT for 72 hrs (n = 3 experiments). B. RT-qPCR analysis showing the effect of Notch signaling inhibition on downstream targets and SOX9. MaSCs derived organoids were treated the same way as for (A) above (n=2 experiments). C. WB analysis showing the effect of NICD1 overexpression on SOX9 in MCF-7 cells (representative results of 4 experiments). D. RT-qPCR analysis showing the effect of NICD1 overexpression on SOX9 and a downstream target, Hey1 (representative results of 4 experiments). E. WB analysis showing the effect of NICD1 overexpression on SOX9 in TNBC cell lines, MDA-MB-231, MDA-MB-436 and SUM159 (representative results of 3 experiments). F. RT-qPCR analysis showing the effect of NICD1 overexpression on downstream targets and SOX9 in TNBC cell lines (representative results of 2 experiments).

### Loss of p53 activates the Notch signaling in mammary organoids

Given that p53 suppresses SOX9 expression in mammary organoids and Notch regulates SOX9 in general, we next asked whether p53 controls Notch signaling and thereby SOX9 expression. We first compared the Notch1 mRNA and Notch1 protein among p53-WT or p53-mut tumor samples in the METABRIC and TCGA datasets (**Fig 3A** and **3B**). We found that Notch1 expression is significantly higher in p53-mutant breast cancer. Experimentally, we analyzed the level of Notch signaling activation between p53-WT and p53-KO MaSC organoids as well as adherent culture. In organoid culture, we could only see a modest increase in NICD1 levels upon p53 loss similar to our observation of the modest effect on SOX9 in organoid condition. However, p53-KO MaSCs had substantially higher levels of NICD1 compared to the control WT organoids when grown in adherent condition (**Fig 3C**). Next, we assessed the expression levels of total Notch and NICD1 in p53-activated cells by Nut-3 treatment. MaSCs treated with Nut-3 led to decreased expression of total Notch, NICD1 and SOX9 altogether (**Fig 3D**). As expected, the decrease of total Notch, NICD1 and SOX9 was only seen in p53-WT organoids upon Nut-3 treatment but not in p53-KO organoids (**Fig 3D, 3E**). Altogether, our results suggest that p53 suppresses SOX9 expression by inhibiting Notch signaling in MaSC organoids.

**Figure 3.**
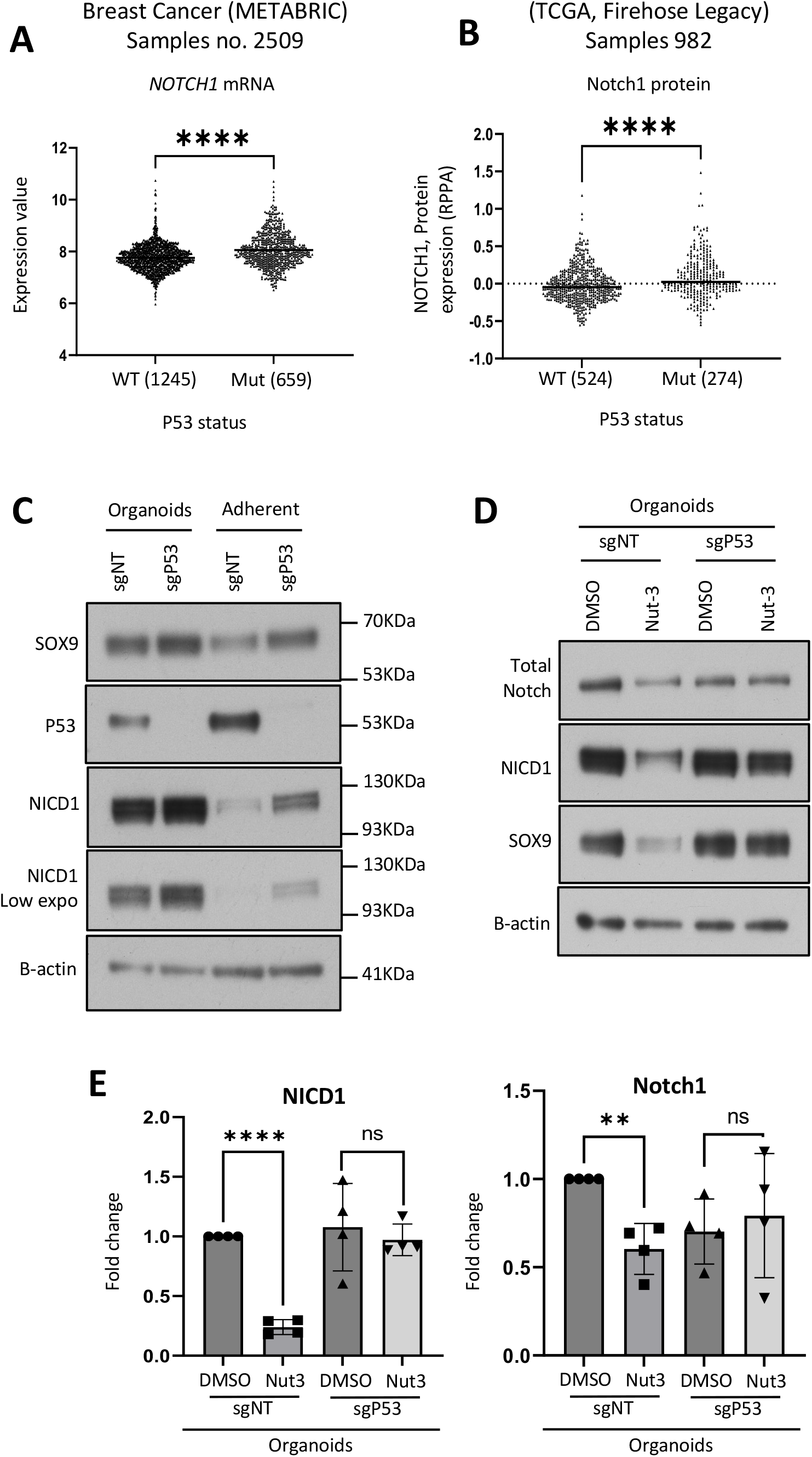
Loss of p53 activates the Notch signaling. **Loss of p53 activates Notch signaling** A. Scatter plots showing the expression of *NOTCH1* mRNA among patients categorized into two groups based on the TP53 status. Analysis performed on METABRIC (n=2509) dataset. B. Scatter plots showing expression of NOTCH1 protein among patients categorized into two groups based on the TP53 status. Analysis performed on TCGA (n=982) dataset. C. WB analysis showing the effect of p53 KO on NICD1 and SOX9 (n=6). Organoids were cultured in organoid condition (3D) and adherent condition (2D) in same media with and without 5% Matrigel, respectively. For 3D culture, 300K cells/well of poly-HEMA coated plates were seeded whereas for 2D culture, 150K cells were seeded in 600mm dishes. MaSCs were grown for 3.5 days before isolating proteins and WB. D. WB analysis showing the effect of p53 activation by Nut-3 treatment on NICD1 and SOX9. MaSC organoids were treated with DMSO or 10uM Nut-3 for 48hrs. E. Quantification of the Notch and NICD1 WB results shown in Fig 3D (n = 4 experiments).

### p53 loss activates Notch signaling via upregulating Psen2

Next, we wanted to understand the mechanism by which p53 controls Notch activation. Notch signaling is activated when the Notch ligand binds to the extracellular domain of the Notch receptor leading to a conformational change in the intracellular domain of the same receptor. Upon sensing the ligand binding, γ-secretase complex then cleaves the intracellular domain, releasing it free from the cell surface, which then induces its targets ^85, 86^. We tested several possibilities, including that p53 loss induces the expression levels of some components of γ-secretase complex or Notch ligands, or p53 loss stabilizes the cleaved intracellular domain of Notch1 to sustain the already activated Notch signaling.

To understand the first possibility, we initially determined the mRNA expression levels of the components of γ-secretase complex (*Psen1, Psen2, Nicastrin* and *Psenen*) upon either p53-KO or Nut-3 treatment (**Fig 4A, 4B, S3A** and **S3B**). Among all, Psen1 and Psen2 are the core catalytic components of the complex. We observed that p53-KO induces the mRNA transcript of *Psen2* (**Fig 4A**), whereas Nut-3 reduces the expression of *Psen2* transcript specifically (**Fig 4C**), suggesting Notch1 signaling activation in p53-KO organoids is probably due to higher levels of Psen2 expression. Databases search identified the presence of multiple TP53-binding sites within the promoter region (−1000 to 100bp relative to transcription start site) of *Psen2* (**Fig S3C)**, further supporting a probable TP53 regulation of Psen2 ^87, 88^. Knowing the fact that only cleaved fragments of either Psen2 or Psen1 forms the γ-secretase complex not the whole protein, we next wanted to analyze the C-terminal fragment (cleaved active fragment) of Psen2 in both p53-WT and p53-KO organoid cells. As expected, we noticed a very evident increase of the CTF-Psen2 protein fragment upon p53 KO and, conversely, a decrease of CTF-Psen2 in Nut-3 treated cells (**Fig 4B** and **Fig 4D**). We have also analyzed the expression levels of the various Notch ligands by RT-qPCR approach and observed an increased expression of only Jag1 upon p53 loss, but this observation was consistent only in organoid cultures but not in adherent cultures (**Fig S3D**), suggesting a possibility of p53-mediated regulation of Notch ligand Jag1 expression which needs further exploration.

**Figure 4.**
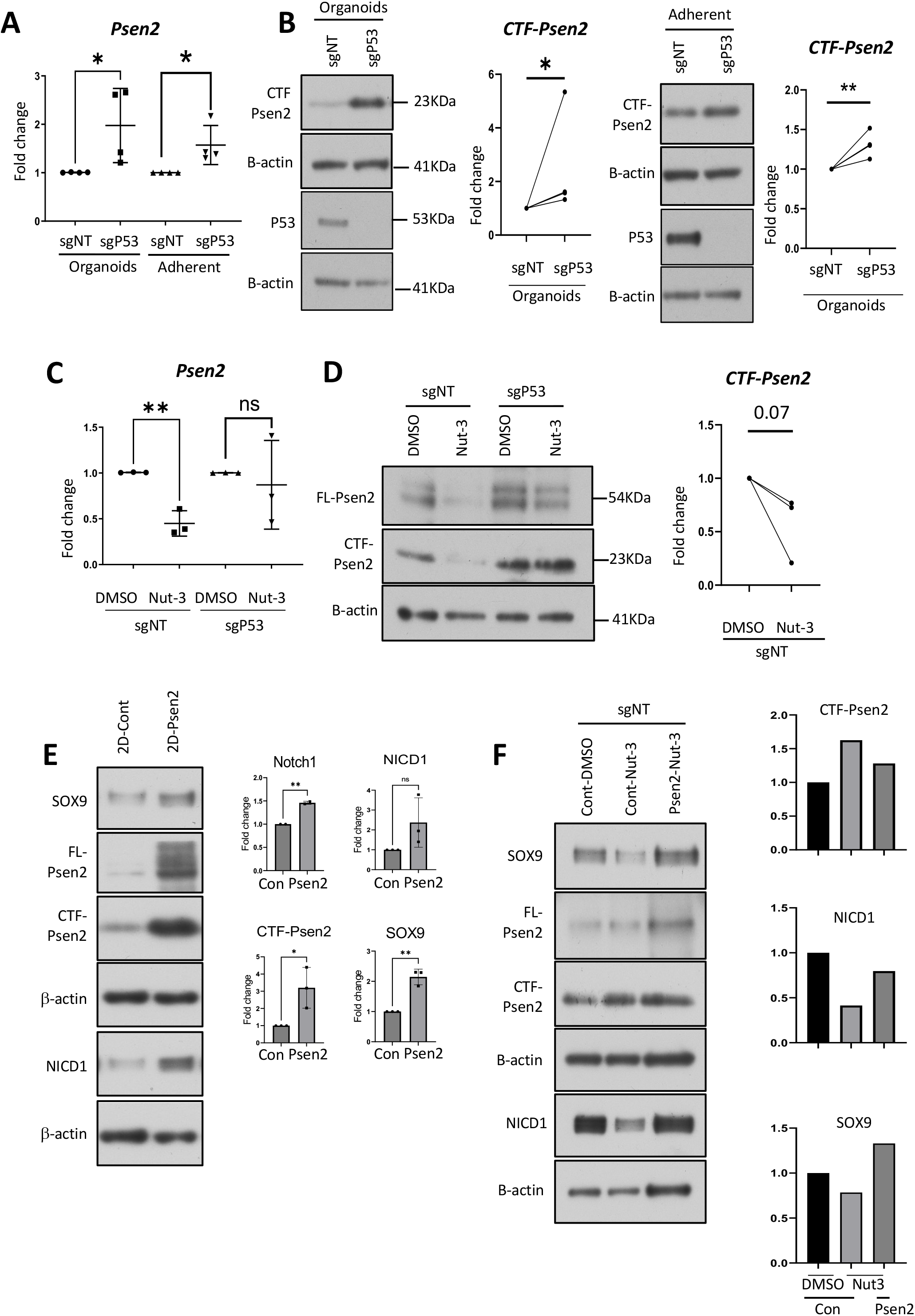
Loss of p53 activates Notch signaling via upregulating Psen2. **p53 controls Notch signaling by inhibiting Psen2 expression** A. RT-qPCR analysis of *Psen2* in the p53-WT and p53-KO organoids (n=4 experiments). B. WB analysis of Psen2 in the p53-WT and p53-KO organoids (n=3 experiments). CTF-Psen2 refers to the C-terminal fragment of Psen2. The graphs show the summarized results of three experiments. C. RT-qPCR analysis of *Psen2* upon p53 activation by Nut-3 treatment in sgNT and sgP53 organoids (n=3 experiments). p53-WT and p53-KO organoids were treated with DMSO and 10uM Nut-3 for 72hrs in organoid culture condition. After 72hrs of treatment, organoids were collected, pelleted and lysed in RNA lysis buffer and isolated RNA was used for RT-PCR. D. WB analysis of Psen2 upon p53 activation by Nut-3 treatment in sgNT and sgP53 organoids (n=3). p53-WT and p53-KO organoids were treated with DMSO and 10uM Nut-3 for 72hrs in organoid culture condition. FL-Psen2 refers to full length Psen2 protein whereas CTF-Psen2 refers to C-terminal fragment of Psen2. Figure 3D and 4D share the same β-actin control blot. Right panel shows the summarized results of three experiments. E. WB analysis showing the effect of Psen2 overexpression on SOX9 in p53-WT organoids (one of two repeats was shown). Organoid cells were grown for 3.5 days in adherent condition, then processed for WB. Right panel shows the quantified CTF-Psen2, SOX9 and NICD1 bands in 2 or 3 experiments. F. WB analysis showing the effect of Nut-3 on SOX9, NICD1 and Psen2 in control and Psen2-overexpressing organoids. Right panel shows the quantified CTF-Psen2, SOX9 and NICD1 bands.

Next, we check the effect of Psen2 overexpression on Notch and SOX9. Psen2-overexpressing organoids, when grown in adherent culture, showed higher expression of both NICD1 and SOX9 compared to the vector control cells (**Fig 4E**). To further connect p53 with Psen2-mediated SOX9 regulation, we increased the levels of p53 by Nut-3 treatment in the normal and Psen2-overexpressing organoids. We noticed that Psen2-overexpressing organoids rescued the levels of NICD1 as well as SOX9 in Nut-3 treated cells (**Fig 4F**)

### p53 loss increases the half-life of SOX9 protein

Next, we asked whether p53 regulation of SOX9 is only mediated by Notch. We treated p53-KO organoids with DAPT in the organoid culture condition. As expected, p53-KO led to higher levels of both NICD1 and SOX9 (**Fig 5A, 5B**). While DAPT fully inhibited NICD1 formation, it only partially downregulated SOX9 in p53-KO cells. These results suggest that p53 could regulate SOX9 through other mechanisms in addition to Notch signaling. Given p53-KO increased SOX9 protein but not transcript levels (**Fig 1**), we asked whether p53 could have any role in the stability of SOX9. To check this notion, we assessed the stability of SOX9 between p53-WT and p53-KO organoids by cycloheximide treatment assay. Surprisingly, we discovered that p53 loss increases the half-life of SOX9 in all our experiments performed with various organoid lines (**Fig 5C, 5D and 5E**). To further test this role, we assessed the stability of SOX9 in MaSCs upon p53 activation by Nut-3 treatment. Consistently, we found Nut-3 decreased SOX9 half-life, supporting that p53 has a regulatory role on SOX9 by modulating its stability (**Fig 5F, 5G**).

**Figure 5.**
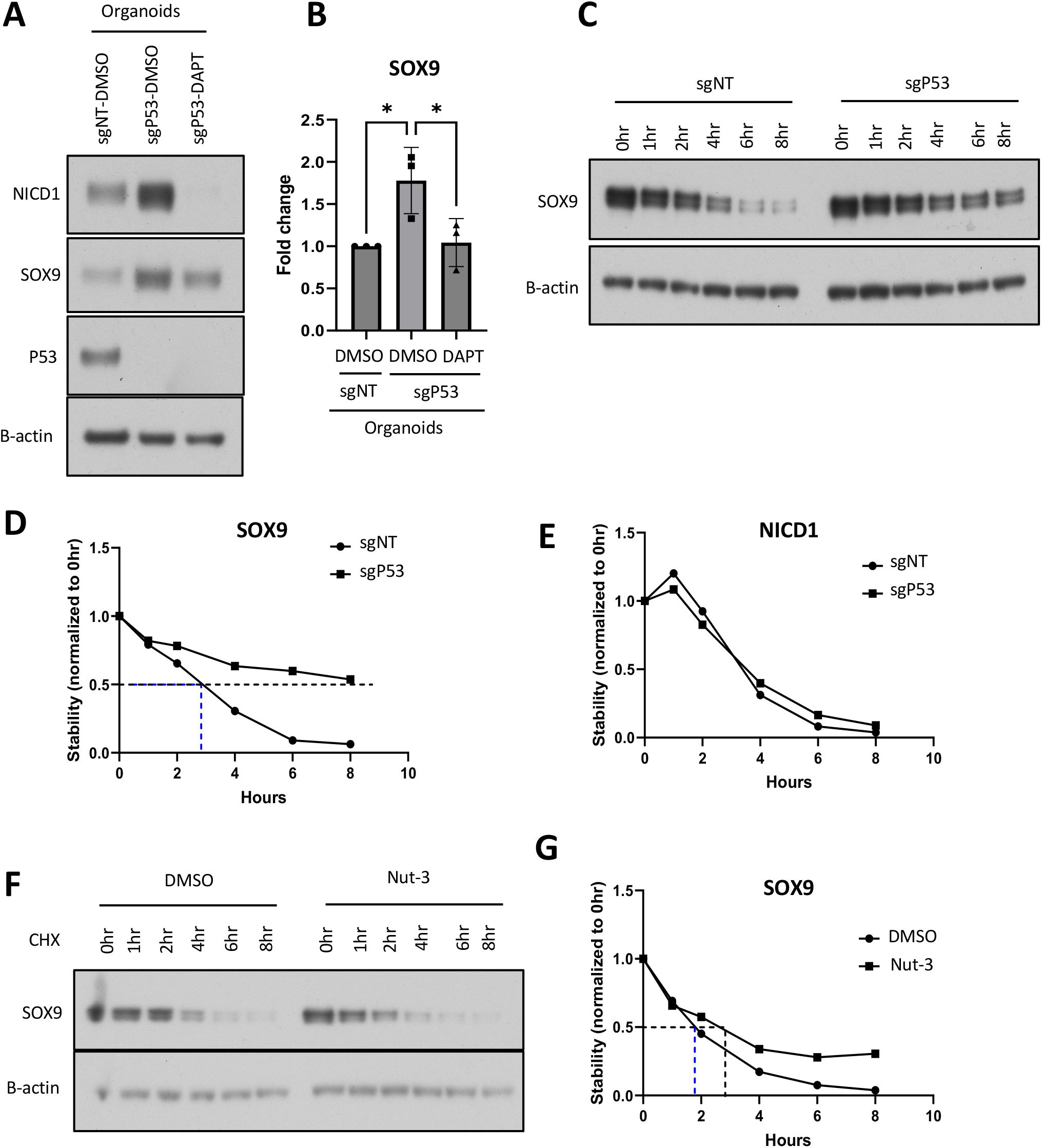
Loss of p53 increases SOX9 protein stability. **P53 reduces SOX9 stability in addition to controlling NOTCH signaling** A. WB showing the effect of DAPT treatment on NICD1 and SOX9 in the sgP53 organoids. 1×10^6 sgP53 organoids were treated with 10uM DAPT or DMSO for 72hrs in organoid culture condition. sgNT organoids treated with DMSO were used as a control. B. Summarized result of SOX9 WB in (A) (n = 3 experiments). C. WB result showing the increased stability of SOX9 in p53-KO organoids. sgNT and sgP53 organoids were treated with 50ug/ml cycloheximide, and the proteins were isolated at different time points and measured for SOX9 by WB. D. Quantification of the SOX9 WB in (C). Stability curve showing the half-life of SOX9 increases more than 2-fold upon p53 loss (representative result of 4 experiments). E. Effect of P53 loss on the stability of NICD1 at different time points. P53 loss does not significantly impact NICD1 stability. F. WB result showing the stability of SOX9 decreases upon p53 activation by Nut-3 treatment. 600K p53-WT organoid cells were treated with DMSO or Nut-3 (10uM) for 48hrs followed by no or added with cycloheximide (50ug/ml) and cultured in organoid condition. Proteins were isolated at different time points and ran in gel. G. Stability curve showing the half-life of SOX9 decreases upon p53 activation by Nut-3 treatment.

## Discussion

Although considerable progress has been made in elucidating the role of SOX9 in driving lineage plasticity along the mammary epithelial cell hierarchy ^6, 27, 28^, how Sox9 is regulated remains an unanswered question in the mammary gland biology and breast cancer field. Among the many recurrent genomic aberrations identified in BLBC, mutation or loss of TP53 is among the most common driver mutations. In the present study, utilizing MaSCs organoid models, we investigated the role of p53 in controlling Sox9 expression. We identified a molecular mechanism by which loss of TP53 activates Notch signaling, which induces Sox9 expression. We have also shown that loss of p53 stabilizes SOX9 protein by almost doubling the half-life of SOX9 protein. We uncovered that the catalytic component of the Notch signaling, Psen2, is upregulated by p53 loss. Based on the literature we know that NICD1 could regulate SOX9 by binding to its promoter ^81, 82, 89^ but whether Hey1 could modulate SOX9 expression is still not known and needs further investigation. Among the Notch ligands, Jag1 appears to be the only one well correlated with the status of TP53 in the RNAs isolated from the organoids grown in native 3D cultures. Whether loss of p53 regulates the SOX9 expression in mammary stem cells in an autocrine manner by producing Jag1 remains to be determined. Alternatively, the stromal compartment can also provide Notch ligands. We believe that in human BLBC patients, SOX9 high expression could be due to the loss of P53 function which drives BLBC progression. The Notch-targeting approach could be effective to suppress the SOX9 expression, thereby limiting BLBC progression.

## MATERIALS AND METHODS

### Cell lines and culture conditions

MCF-7, MDA-MB-231, MDA-MB-436, SUM159, were cultured in DMEM supplemented with 10% FBS + 1% Pen-Strep antibiotics. HEK293T cells were cultured in DMEM supplemented with 10% FBS supplemented with 1% sodium pyruvate and 1% NEAA (non-essential amino acids). All the cell lines were grown in humidified incubator containing 5% CO_2_ and maintained at 37°C.

### Mouse models

Mice from FVB background expressing SOX9-GFP and C(3);Tag/SOX9-GFP were used for the study. The age of the mice used for the study was between 4 weeks to 8 weeks.

### Immunoblotting

For 2D cultured cells, media was aspirated out and the cells were rinsed with 1xPBS. RIPA buffer was directly added to the monolayer, cells were scraped with cell scrapper and collected in tube. Lysates were further cleared by passing the lysates through syringes and incubated in ice for a minimum of 30 minutes. After 30mnts lysates were centrifuged at 14000 rpm for 15 min and proteins were collected and stored in-80°C or immediately used for WB. For organoid cultures, organoids were collected and Matrigel was broken/diluted using ice-cold PBS and centrifuged at 500g for 5 min. The obtained organoid pellets were directly lysed in RIPA lysis buffer with the help of syringe. Organoid lysates were centrifuged at 14000 rpm for 15 min and the collected proteins were run in the precast Bis-Tris protein gels 4-12%. Resolved proteins were transferred to PVDF membrane, blocked with 5% non-fat milk for 1hr, washed with 1xPBST three times, 10min each and incubated with primary antibodies overnight in cold room. Next day, the blots were taken out from the primary antibodies and washed thrice with 1xPBST 10min each and incubated with secondary antibodies for 1hr at RT, washed thrice with 1xPBST 10min each and developed using ECL reagent in darkroom.

### RT-qPCR

For 2D cultured cells, media was aspirated out and the cells were rinsed with 1xPBS. RNA lysis buffer was directly added to the monolayer, cells were pipetted up and down and collected in tube. For organoid cultures, organoids were collected and Matrigel was broken/diluted using ice-cold PBS and centrifuged at 500g for 5 min. The obtained organoid pellets were directly lysed in RNA lysis buffer. RNAs were isolated from the lysed cells using RNA isolation kit as per the manufacturer’s protocol. The concentration and the purity of the isolated RNAs were checked using nanodrop based on 260/280 and 260/230 ratios and converted to cDNA using random hexamer-based cDNA synthesis kit (Applied Biosystems). Synthesized cDNAs were further diluted with nuclease free water, if required and used for RT-qPCR analysis of the required genes. For qRT-PCR experiment, QuantStudio 6 software (Applied Biosystems) was used.

### DAPT treatment

MaSCs derived organoids were trypsinized and live cells were counted using trypan blue. 300K live cells were mixed with 2ml organoid media with a final conc. of 10uM DAPT and 5% Matrigel and seeded in one well of poly-HEMA coated 6-well plates. After 72hrs of treatment, organoids were collected, proteins and RNAs were isolated, WB and RT-PCR were performed.

### Nutlin-3 treatment

p53-WT and p53-KO organoids were treated with DMSO control and 10uM Nut-3 for 48hrs in organoid culture condition. Briefly, organoids were trypsinized and live cells were counted using trypan blue. 400K live MaSCs were mixed with 2ml organoid media with a final conc. of 10uM Nut-3 and 5% Matrigel and seeded in one well of poly-HEMA coated 6-well plates. After 48hrs of treatment, organoids were collected and Matrigel was broken/diluted using ice-cold PBS and centrifuged at 500g for 5 min. The obtained organoid pellets were directly lysed in RIPA lysis buffer with the help of syringe. Organoid lysates were centrifuged at 14000 rpm for 15 min and the collected proteins were run in the precast Bis-Tris protein gels 4-12%. Resolved proteins were transferred to PVDF membrane, blocked with 5% non-fat milk for 1hr, washed with 1xPBST three times, 10min each and incubated with primary antibodies overnight in cold room. Next day, the blots were taken out from the primary antibodies and washed thrice with 1xPBST 10min each and incubated with secondary antibodies for 1hr at RT, washed thrice with 1xPBST 10min each and developed using ECL reagent in darkroom.

### Mammary gland isolation and cell preparation

Mammary glands were collected in sterile tube containing sterile 1xPBS, examined under microscope and minced with razor blade to 1mm fragment size. Minced glands were kept for digestion for 1.5-2hrs in 37°C in AdDMEM media containing collagenase, dispase and RI in 15ml tube. After digestion was completed, 1xPBS was added till 10ml and centrifuged at 500g for 5min. The top fatty layer was aspirated out and the pellet was treated with 1ml RBC lysis buffer to lyse red blood cells. After 1 min incubation, reaction was halted by adding 1xPBS on top till 10ml, centrifuged at 500g for 5 min. Pellet was mixed with 0.05% Trypsin and incubated in 37°C water bath for 5 min, after which the trypsinization process was quenched by adding Ad/DMEM media containing 10% FBS, centrifuged and pelleted down. To the pellet 1ml media Dispase and DNAse was added and incubated in 37°C water bath for 5 min. Reaction was halted by adding 1xPBS on top till 10ml and centrifuged at 500g for 5min.

### Organoid Culture

Fresh isolated MaSCs from mice breast tissue were resuspended in ice cold Epicult-B media + 5% FBS + 10 ng/ml EGF + 20 ng/ml FGF2 + 4 mg/ml Heparin + 5 μM Y-27632 + 5% Matrigel (Corning 354234). This mixture was then seeded in 2-hydroxyethyl methacrylate (Poly-HEMA) coated 6-well plates (300K-500K cells /well). For organoid passaging, organoids were collected, washed with PBS, disassociated with 0.05% Trypsin-EDTA and then reseeded in organoid culture medium. For regular passaging MaSCs were maintained in a seeding density 300K cell/well. Cells were passaged every 3-4 days depending on the densities of the particular genetically modified cell type.

### CRISPR sgRNA Design and Cloning

All the sgRNA sequences used in the paper including p53, Psen2 sgRNAs were designed using the CHOPCHOP webserver (Labun et al., 2016). Oligos were cloned into pLenti-CRISPRv2.Puro and pLentiCRISPRv2.Blast vectors. The sgRNA targeting sequences are in Table S2.

### Transformation and plasmid preparation

Cloned plasmids were transformed into ElectroMAX Stbl4 Competent Cells (ThermoFisher 11635018) using a MicroPulser Electroporator (BIO-RAD) using cuvettes with a 1 mm gap and the default settings for E. coli. Plasmids were isolated from 150ml bacterial cultures using ZymoPURE™ II Plasmid Purification Kit, Maxiprep Cat # D4203.

### Lentivirus generation

For virus particles production, lenti-packaging (pCMV-deltaR8.74) and VSVG (pMD2.G) plasmids were used along with the Lentiviral based vectors either for overexpressing or knocking out a gene. Virus production was performed on collagen coated 100mm culture dishes.

Briefly, the plates were coated with 2ml of 10-20µg/ml collagen I (Corning #354236) solution prepared in sterile 1xPBS (Fisher Scientific #SH3025601) and incubated for 1hr in tissue culture hood. After 1hr, HEK293T cells were trypsinized and seeded to a cell density of ∼70%. Next day, the plates were transfected with following plasmids-pCMV-deltaR8.74, pMD2.G, and respective vector in the following ratio of 1.8µg:1.2µg:3µg respectively, in the HEK293T media i.e., 10ml DMEM (Corning #10-017-CV) supplemented with final concentrations of 10% FBS (Corning™ 35016CV), MEM 1xNEAA (Gibco #11140-050) and 1mM Sodium pyruvate (Gibco #11360-070). For transfection, the plasmids were diluted in 600µl jetPRIME® Buffer (Polyplus #712-60/60ml) and mixed. 12µl of jetPRIME® transfection reagent (Polyplus #101000046) was added to this mixture, vortexed briefly, spun briefly and incubated for 10min at RT. The mixture was overlayed on the HEK293T cells cultured overnight in 100mm dishes after 10min of incubation and the dishes were swirled gently. Post 6hrs of transfection the media was aspirated out and replaced with fresh HEK293T culture media. Viral sups were collected at 24hrs, 48hrs and 72hrs time points and replaced with fresh media each time. At the end of 72hrs, collected viruses at different time points were pooled together, concentrated to 1/20-fold to the original volume using Lenti-X Concentrator (Clontech Laboratory #631232) and stored in-80°C.

### CRISPR mediated organoid genetic manipulation

MaSCs were infected with a MOI of 1. Briefly 300K cells were infected with viruses and seeded in Poly-HEMA) coated 6-well plates for 24hrs. Next day cells were collected in 15ml tube and washed with sterile ice-cold 1xPBS once, reseeded in the wells with fresh organoid media for another 24hrs. Next day antibiotic selection was begun depending on the marker present in backbone of the lentivirus vector used for cloning. Once the cells were completely selected, cells were always maintained in half dose of the selection antibiotic. Knockout efficiencies were checked by percentage of indel via TIDE assay first followed by RT-PCR and/or WB, whereas overexpression was verified by RT-PCR and/or WB in the tested cells. Briefly genomic DNA was extracted from the cells, and 100 ng of genomic DNA was used in a PCR amplification using SapphireAMP Fast PCR Master Mix (Clontech Cat# RR350B) and appropriate primers surrounding the sgRNA target region. An amplified PCR product was sent for sequencing, and the sequencing results were checked for indel efficiencies using TIDE assay.

### Ligands treatment

Treatments were given in 6-well plate. Briefly, 600K cells/well were seeded in 4ml Ad/DMEM media containing 10% FBS with all four growth factors. Cells were cultured for 3 days before giving treatments. Ligands were treated at the final concentration of 100ng/ml for Wnt3a, 500ng/ml for R-spondin; 100ng/ml for FGF10 and 5ng/ml or 10ng/ml for TGFb. Treatments were given for 48hrs or 72hrs. For FGF10 treatment, FGF2 was replaced with FGF10 in the final media.

### Hypoxia treatment

Organoids were trypsinized, counted and 600K cells/well were seeded in 4ml Ad/DMEM media containing 10% FBS with all four growth factors. Cells were cultured for 3 days in humidified incubator containing 10% CO_2_ and maintained at 37°C. The plates were then moved either in the normoxia incubator containing 21% oxygen supply and the hypoxia incubator set at 1% oxygen. Treatments were given for 24hrs and 48hrs.

### Bioinformatics analysis

To examine the expression levels of SOX9 transcript among patients with WT P53 and Mut P53, TCGA and METABRIC datasets have been used. To see the expression levels of Notch1 transcript and Notch1 proteins among patients with WT P53 and Mut P53, TCGA and METABRIC datasets have been used. Expression values were extracted, and graphs are made using Graph Pad prism.

## Statistics

Where necessary, statistical data were analyzed using GraphPad Prism to generate SE values and to determine the level of significance using the student’s t-test (one-tailed or two-tailed, as appropriate) and ANOVA; *p-value < 0.05; **p-value 0.005, ***p-value 0.0001 were considered to indicate significance. Data are reported as mean ± SEM. A two-tailed Student’s t-test was used for comparisons of continuous variables between two groups. One-way analysis of variance (ANOVA) was used when three or more groups were compared. A two-way ANOVA with multiple comparisons test was used to check the effect of two factors on a dependent variable. All details regarding n number and what n represents are stated in figure legends.

## Acknowledgments

We thank the Flow Cytometry, Histopathology, Analytical Imaging, Genomics, and Stem Cell Isolation core facilities of Albert Einstein College of Medicine for technical assistance (supported by Albert Einstein Cancer Center support grant P30 CA013330, New York State Department of Health NYSTEM program shared facility grant C029154, and NIH grant 1S10OD019961-01 for the P250 High-Capacity Slide Scanner). This work is supported by the NIH grant 1P01CA257885-01A1 (W.G.).

## Author Contributions

Conceptualization, RS and WG; Methodology, RS and WG; Investigation, RS and WG; Visualization RS and WG; Funding acquisition, W.G. RS and WG conceived the study; R.S performed all the experiments, acquired, analyzed data, and wrote the original draft; all authors contributed to data interpretation; R.S. and W.G. wrote the manuscript.

## Supplemental Figure Legends

**Figure S1.**
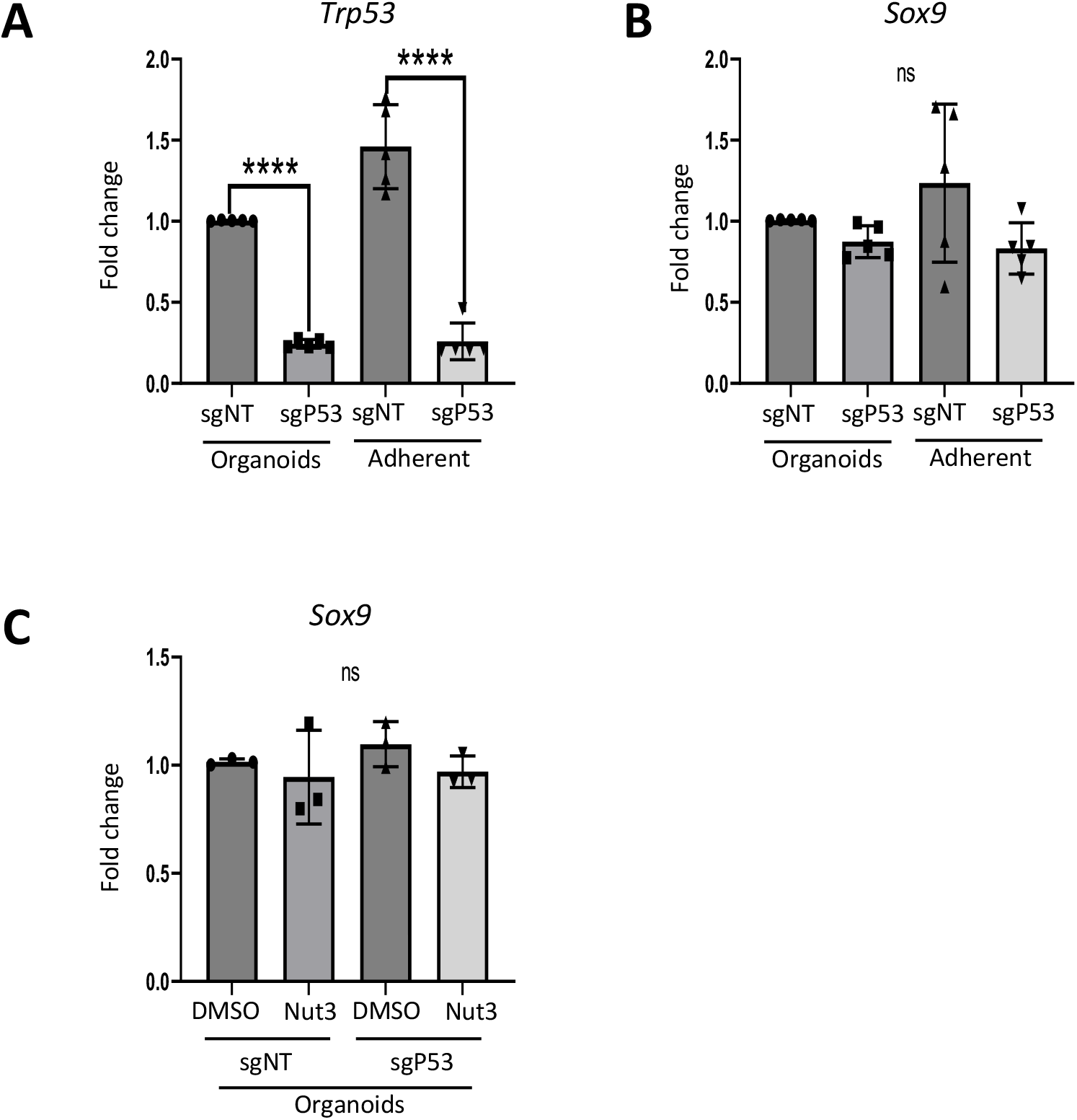
Loss of p53 upregulates SOX9 expression. A) RT-qPCR analysis of *Trp53* in the p53-WT or p53-KO organoids (n=5 experiments). B) RT-qPCR analysis of *Sox9* in the p53-WT or p53-KO organoids (n=5 experiments). C) RT-qPCR analysis of *Sox9* in the sgNT and sgP53 organoids treated with DMSO or 10uM Nut-3 for 48hrs in organoid culture (n=3 experiments).

**Figure S2.**
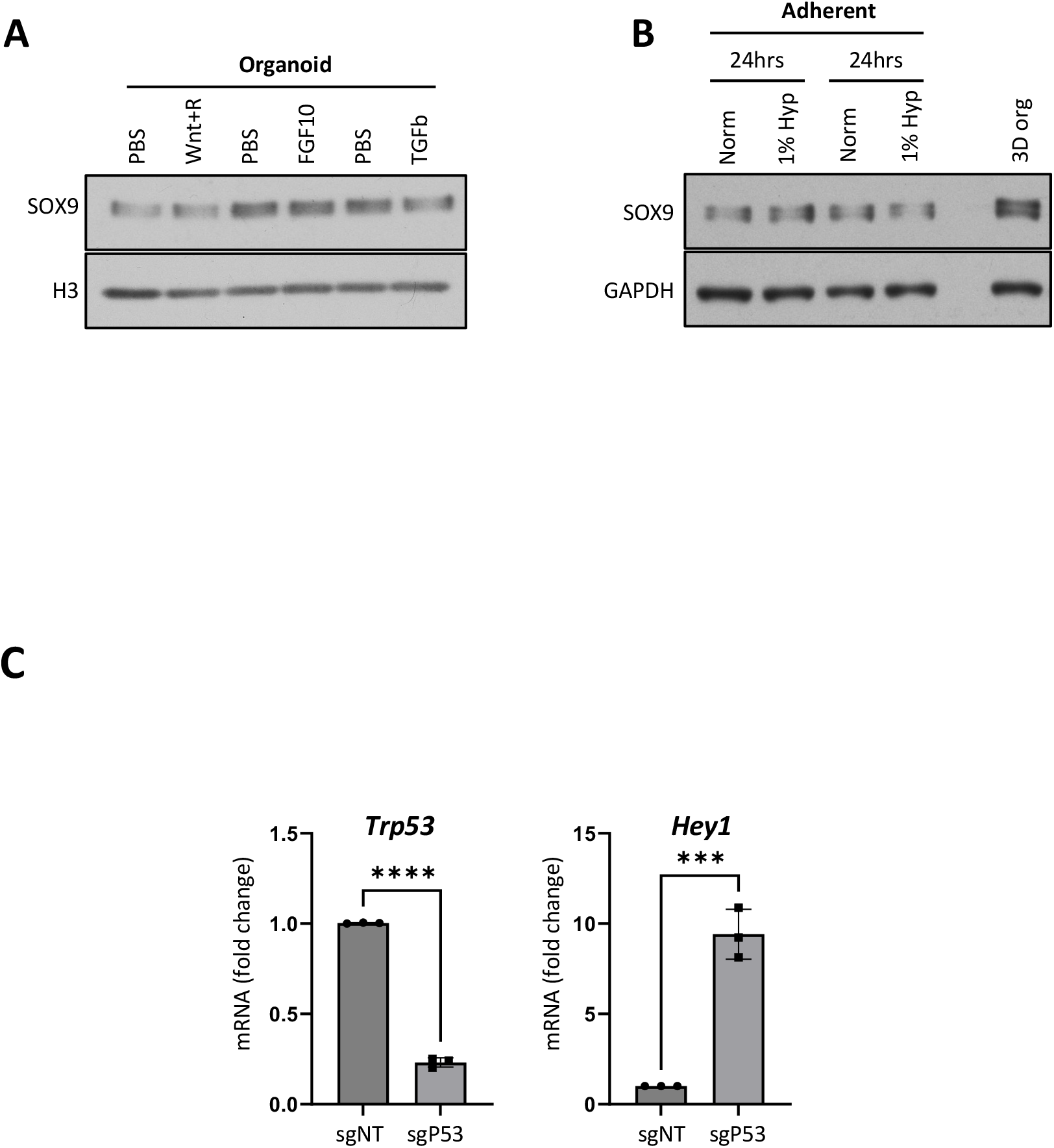
NOTCH signaling upregulates SOX9. A. WB analysis of the effect of Wnt3a, FGF10 and TGFβ ligands treatment on SOX9. 600K cells were cultured in adherent condition in 60mm dishes for 3 days in 2ml organoid culture medium without Matrigel. On third day, media was refreshed with ligands containing media. Ligands were treated at the final concentration of 100ng/ml for Wnt3a, 500ng/ml for R-spondin; 100ng/ml for FGF10 and 5ng/ml for TGFβ. For FGF10 treatment, FGF2 was replaced with FGF10 in the final media. Treatments were given for 48hrs. B. WB analysis of the effect of 1% hypoxia on SOX9. Organoid cells (600K) were cultured in adherent condition in 60mm dishes for 3 days in 2ml organoid culture medium without Matrigel. On third day, media was refreshed, and the cells were cultured in normoxia (21% oxygen) or hypoxia (1% oxygen) for 24hrs or 48hrs. C. RT-PCR analysis of the Hey1 in control vs p53-KO MaSCs. Hey1 is the most consistently observed Notch downstream target affected by p53 status.

**Figure S3.**
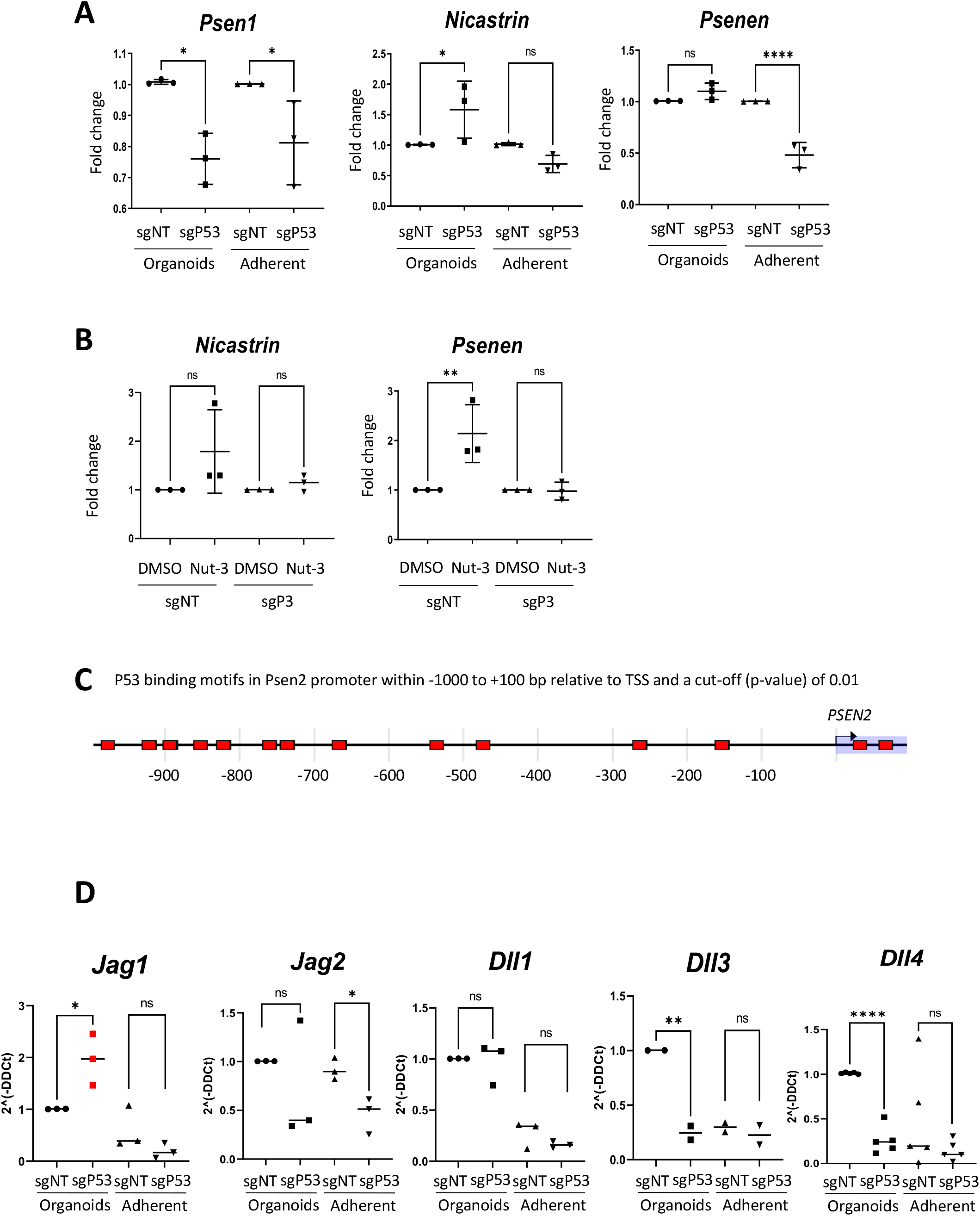
p53 activates Notch signaling via Psen2. **p53 controls Notch signaling by inhibiting Psen2 expression** A) RT-qPCR analysis of *Psen1, Nicastrin and Psenen* in the p53WT and p53KO organoids (n=3 experiments). B) RT-qPCR analysis of *Nicastrin* and *Psenen* upon p53 activation by Nut-3 treatment in sgNT and sgP53 organoids (n=3 experiments). Organoids were treated with DMSO and 10uM Nut-3 for 48hrs in organoid culture condition. C) Diagram showing the presence of putative P53 binding sites in Psen2 promoter within-1000 to +100 bp relative to TSS with a cut-off (p-value) set as 0.01. EPD (Eukaryotic Promoter Database) has been used to identify the binding sites. D) RT-qPCR analysis of Notch ligands, *Jag1, Jag2, Dll1, Dll3, and Dll4*, in the p53-WT and p53-KO organoids cultured in the same way as in (A) (n=3 experiments).

