## Supplemental Table for "Loss of tumor suppressor p53 upregulates stem cell factor SOX9 via Notch signaling"

**Table S1: Reagent or resources**

| **Reagent or resource** | **Source** | **Identifier** |
| --- | --- | --- |
| **Antibodies** |  |  |
| Biotin anti-mouse TER-119 | BioLegend | AB_313705 |
| Biotin anti-mouse CD45 | BioLegend | AB_312969 |
| Biotin anti-mouse CD31 | BioLegend | AB_312899 |
| PerCP/Cy5.5 anti-mouse/human CD49f | BD Biosciences | AB_11151910 |
| APC anti-mouse CD326 (Ep-CAM) | BioLegend | AB_1134102 |
| PE/Cy7 anti-mouse Ly-6A/E (Sca-1) | BioLegend | AB_493596 |
| StreptavidinV450 | BD Biosciences | AB_2033992 |
| Sox9 (D8G8H) Rabbit mAb | Cell Signaling Technology | AB_2665492 |
| Rabbit Anti-Histone H3 | Abcam | AB_302613 |
| Mouse anti-β-actin | BD Biosciences | 612656 |
| Mouse anti-TrP53 | Cell Signaling | 2524S |
| Rabbit anti-CTF-Psen2 | Cell Signaling | 9979 |
| Rabbit anti-NICD1 | Cell Signaling | 4147S |
| HRP Donkey anti-rabbit IgG | BioLegend | 406401 |
| HRP Donkey anti-mouse IgG | BioLegend | 405306 |
| Goat anti-Rabbit IgG (H+L) Cross-Adsorbed Secondary Antibody, Alexa Fluor 647 | Thermo Fisher | AB_2535812 |
| **Bacterial and Virus Strains** |  |  |
| ElectroMAX Stbl4 Competent Cells | Thermo Fisher | Cat#11635018 |
| **Chemicals, Peptides, and Recombinant Proteins** |  |  |
| DAPT | Cayman Chemical Company | 13197 |
| Nutlin-3 | Cayman Chemical Company | 10004372 |
| Sodium Chloride | Fisher Scientific | BP358212 |
| Ethanol | Decon Laboratories, Inc. | 2701 |
| Chloroform | Fisher Scientific | BP1145 |
| Glacial Acetic Acid | Fisher Scientific | A38S500 |
| 1M Tris-HCl, pH8.0 | TekNova | T1080 |
| Magnesium Chloride Hexahydrate | Fisher Scientific | BP214-500 |
| Calcium Chloride | Fisher Scientific | BP510-500 |
| Nonidet P-40 Substitute | IBI Scientific | IB01140 |
| Sodium Deoxycholate | Sigma-Aldrich | 30970 |
| Sodium Orthovanadate | Sigma-Aldrich | S6508 |
| Sodium Dodecyl Sulfate | Fisher Scientific | BP-166 |
| 10% Formalin | Fisher Scientific | SF98-4 |
| Esp3I (BsmBI) | New England Biolabs | R0734S |
| BamHI-HF® Restriction Enzyme, 20,000 units/ml, 10,000 units | New England Biolabs | R3136S |
| T4 DNA Ligase | New England Biolabs | M0202S |
| Jetprime transfection reagent | VWR | 89129-924 |
| Advanced DMEM/F12 medium | Life Technology | 12634010 |
| Epidermal Growth Factor human (EGF) | Sigma-Aldrich | E9644 |
| Fibroblast Growth Factor basic Protein, Human recombinant (FGF2) | EMD Millipore | GF003 |
| Heparin sodium salt, from porcine intestinal mucosa | EMD Millipore | H3149 |
| Y-27632 (hydrochloride), ROCK inhibitor | Cayman Chemical Company | 10005583-10 |
| Matrigel | Corning | 354234 |
| Red blood cell lysing buffer | Sigma-Aldrich | R7757 |
| EpiCultTM-B Mouse Medium | Stem Cell Technologies | 05610 |
| Poly(2-hydroxyethyl methacrylate) | Sigma-Aldrich | P3932-25G |
| Ethanol 200 Proof | Becon Labs | 2701 |
| Phosphate-Buffered Saline, 1X without calcium and magnesium, pH7.4±0.1 | Corning | 21-040-CM |
| Trypsin 0.05% with 0.53 mM EDTA | Corning | 25-052-CI |
| Dimethylsulfoxide (DMSO) | Millpore-Sigma | MX-1458 |
| Carbenicillin (Disodium) | Goldbio | C-103-5 |
| Puromycin dihydrochloride | Tocris | 4089 |
| Blasticidin S HCl | Corning | MT30100RB |
| Polybrene | EMD-Millipore | TR-1003-G |
| Rat-tail collagen I | Corning | 354236 |
| 30% hydrogen peroxide | Sigma-Aldrich | 31642 |
| Antigen unmasking solution, Citric Acid Based | Vector Laboratories | H-3300 |
| Thermo Scientific™ Halt™ Protease Inhibitor Cocktails | Fisher Scientific | PI87786 |
| Thermo Scientific™ Halt™ Phosphatase Inhibitor | Fisher Scientific | PI-78420 |
| BLUEstain™ Protein ladder | Goldbio | P007-3000 |
| NuPAGE® LDS Sample Buffer (4X) | Invitrogen | NP0007 |
| NuPAGE® MOPS SDS Running Buffer (for Bis-Tris Gels only) (20X) | Invitrogen | NP0001 |
| NuPAGE® Transfer Buffer (20X) | Invitrogen | NP00061 |
| Millipore* Immobilon* PVDF Transfer Membranes | Millipore | IPVH00010 |
| Carnation Instant Nonfat Dry Milk | Amazon | B07NQDB3XS |
| BSA fraction V | Fisher Scientific | BP1600100 |
| Western Blot Stripping buffer | Thermo Fisher Scientific | 46430 |
| Western Lightning ECL Pro | PerkinElmer | NEL121001EA |
| DMEM/F12 medium | Corning | 10-092-CV |
| Deoxyribonuclease I (DNase I) | Worthington | LS002139 |
| Hyaluronidase | Worthington | LS002592 |
| Collagenase, Type III | Worthington | LS004182 |
| Neutral Protease (Dispase) | Worthington | LS02109 |
| RBC lysis buffer | eBioscience | 00-4300-54 |
| UltraPure TM 0.5M EDTA, pH8.0 | Thermo Fisher | 15575020 |
| Dulbecco’s Modified Eagle Medium (DMEM) | Corning | 10-017-CV |
| RPMI1640 | Gibco | 11875085 |
| Fetal Bovine Serum, Heat Inactivated | Corning | 35-015-CV |
| MEM Non-Essential Amino Acids Solution 100mM (100X) | Invitrogen | 11140-050 |
| MEM Sodium Pyruvate Solution 100mM (100X) | Invitrogen | 11360-070 |
| GlutaMAX™ Supplement 100x | Gibco | 35050061 |
| Penicillin-streptomycin 100x | Corning | 30-002-CI |
| Retronectin | Takara | T100A |
| β-mercaptoethanol | Sigma-Aldrich | M6250-100ML |
| 32% Paraformaldehyde (Formaldehyde) aqueous | Electron Microscopy Sciences | 15714-S |
| TritonX-100 | Sigma-Aldrich | BP151 |
| Tween 20, molecular biology grade | Sigma-Aldrich | X100 |
| Xylene | Fisher Chemical | BPX3P1GAL |
| DAPI | Sigma-Aldrich | D8417 |
| DAPI Fluoromount-G mounting medium | Southernbiotech | 0100-20 |
| Fisher Chemical™ Permount™ Mounting Medium | Fisher Scientific | SP15100 |
| Trizol LS | Thermo Fisher | 10296010 |
| Methanol | EMD Millipore | MX0488-1 |
| Isopropanol | Fisher Scientific | BP26184 |
| Lenti-XConcentrator | Takara | 631232 |
| VisiGlo Prime HRP Chemiluminescent ECL Substrate | Amresco | 89424-016 |
| **Critical Commercial Assays** |  |  |
| Quick Ligation Kit | New England Biolabs (NEB) | M2200S |
| High-Capacity cDNA Reverse Transcription Kit | Applied Biosystems | 4368814 |
| SapphireAmp® Fast PCR Master Mix | Clontech | RR350B |
| Power SYBR® Green PCR Master Mix | Applied Biosystems | 4368708 |
| Direct-zol RNA Miniprep Plus kit | Zymo Research | R2050S |
| Zyppy™ Plasmid Miniprep Kit | Zymo Research | D4020 |
| ZymoPURE II Plasmid Maxiprep Kit | Zymo Research | D4203 |
| NucleoSpin® Gel and PCR CleanUp | Takara | 740609 |
| PrimeSTAR MAX DNA polymerase | Takara | R045A |
| DAB Peroxidase (HRP) Substrate Kit, 3,3’-diaminobenzidine | Vector Laboratories | SK-4100 |
| Dura Chemiluminescent Substrate | Fisher Scientific | PIA34075 |
| NuPAGE™ Novex™ 4-12% BisTris Protein Gels, 1.0 mm, 12-well | Invitrogen | NP0322BOX |
| **Experimental Models: Cell Lines** |  |  |
| Mouse organoid: Sox9-GFP;C3-TAg mammary epithelial organoid | This paper | N/A |
| Mouse organoid: Sox9-GFP mammary epithelial organoid | This paper | N/A |
| **Experimental models: Organisms/strains** |  |  |
| Mouse: FVB-Tg(C3-1-TAg)cJeg/JegJ | The Jackson Laboratory | JAX:013591; RRID:IMSR_JAX:013591 |
| **Oligonucleotides** |  |  |
| qRT-PCR primers | This paper | Table S2 |
| sgRNA sequences | This paper | Table S2 |
| Genotyping primers | This paper | Table S3 |
| Full length mPsen2 gene | This paper | NCBI |
| **Recombinant DNA** |  |  |
| pLVX-Puro | Takara | 632164 |
| pLVX-NICD1 | This paper | N/A |
| pMD2.G | Addgene | Addgene_12259 |
| pCMVR8.74 | Addgene | Addgene_22036 |
| pLenti-CRISPRv2.Puro | Addgene | Addgene_98290 |
| pLenti-CRISPRv2.Puro-sgNT | This paper | N/A |
| pLenti-CRISPRv2.Puro-sgP53 | This paper | N/A |
| pLV-EF1a-IRES-Blast | This paper | N/A |
| pLV-EF1a-IRES-FL. mPsen2 | This paper | N/A |
| pLenti-CRISPRv2.Blast-sgNT | This paper | N/A |
| pLenti-CRISPRv2.Blast-sgPsen2 | This paper | N/A |
| **Software and algorithms** |  |  |
| QuantStudio 6 Real-Time PCR Software | Applied Biosystem | N/A |
| GraphPad Prism (version 9.3.0) | Dotmatics | <https://www.graphpad.com> |
| ImageJ (FIJI, version 2.9.0) | Schindelin et al. 69 | <https://imagej.net/software/fiji/> |
| BioRender | BioRender | <https://app.biorender.com/> |
| **Other** |  |  |
| Cell strainer, 40 μm | Fisher Scientific | 22-363-547 |
| 0.22 μm pore size sterile syringe filter | Foxx Life Sciences | Cat# 371-2215-OEM |
| Whatman Puradisc 25 mm Polyethersulfone Syringe Filter, 0.45 µm, sterile | Cytiva | 6780 -2504 |
| FILM, HYBLOT, HIGH SENSITIVITY 5x7" | Thomas Scientific | 1156P38 |
| Cell Scrapers | Celltreat | 229311 |
| 1mm electroporation Cuvettes | Thermo Fisher Scientific | FB101 |

**Table S2: RT-PCR Primers and sgRNA sequences used in the study**

| **RT-Primer** | **Sequence** |
| --- | --- |
| mTrp53 RT-Fwd | CTCTCCCCCGCAAAAGAAAAA |
| mTrp53 RT-Rev | CGGAACATCTCGAAGCGTTTA |
| mSox9 RT-Fwd | AGGAAGCTGGCAGACCAGTA |
| mSox9 RT-Rev | CGTTCTTCACCGACTTCCTC |
| mP21 RT-Fwd | TAAGGACGTCCCACTTTGCC |
| m P21 RT-Rev | CGTCTCCGTGACGAAGTCAA |
| mMdm2 RT-Fwd | AAGATGCGCGGGAAGTAGC |
| mMdm2 RT-Rev | GCACCCTCGGTAGACACAG |
| mPuma RT-Fwd | TGTGGATCTGCAGGTGTCTC |
| mPuma RT-Rev | CTAGACCCTCTACGGGCTCC |
| mBax RT-Fwd | CTGGATCCAAGACCAGGGTG |
| mBax RT-Rev | CCTTTCCCCTTCCCCCATTC |
| mFas RT-Fwd | GTCCTGCCTCTGGTGCTTG |
| mFas RT-Rev | AGCAAAATGGGCCTCCTTGA |
| mHes1 RT-Fwd | CCAGCCAGTGTCAACACGA |
| mHes1 RT-Rev | AATGCCGGGAGCTATCTTTCT |
| mHey1 RT-Fwd | CGAGAAGCGCCGACGAGACC |
| mHey1 RT-Rev | CAGGGCGTGCGCGTCAAAAT |
| mPsen1 RT-Fwd | CCCAAAGGCCCACTTCGTAT |
| mPsen1 RT-Rev | GGTACCCTCCTTTGGGCTTC |
| mPsen2 RT-Fwd | TAAACTCTACCCCGCGCC |
| mPsen2 RT-Rev | CCGCTCATCACACACCTCTT |
| mNicastrin RT-Fwd | TTTTCCGTGGTACTGGCAG |
| mNicastrin RT-Rev | CCCCTGTATCCCCACTAATTG |
| mPsenen RT-Fwd | TCTTGGTGGATTTGCGTTCC |
| mPsenen RT-Rev | GAACGCCTCTCTGAAGAACCA |
| mJag1 RT-Fwd |  |
| mJag1 RT-Rev |  |
| mJag2 RT-Fwd | CGTGGCTGCTATCACTCAGA |
| mJag2 RT-Rev | AGCCACAGCACACTGAACAC |
| mDll1 RT-Fwd | GACCTCGCAACAGAAAACCCA |
| mDll1 RT-Rev | TCCGTAGTAGTGCTCGTCACA |
| mDll3 RT-Fwd |  |
| mDll3 RT-Rev |  |
| mDll4 RT-Fwd |  |
| mDll4 RT-Rev |  |
| mGAPDH RT-Fwd | AGGTCGGTGTGAACGGATTTG |
| mGAPDH RT-Rev | TGTAGACCATGTAGTTGAGGTCA |
| hSox9 RT-Fwd | AGTACCCGCACTTGCACAAC |
| hSox9 RT-Rev | CGTTCTTCACCGACTTCCTC |
| hHes1 RT-Fwd | GTGTCAACACGACACCGGAT |
| hHes1 RT-Rev | ATGCCGCGAGCTATCTTTCT |
| hHey1 RT-Fwd | GTGCGGACGAGAATGGAAAC |
| hHey1 RT-Rev | TTGCTCCATTACCTGCTTCTC |
| hGAPDH RT-Fwd | CGGTTTCTATAAATTGAGCCCGCAG |
| hGAPDH RT-Rev | ACCAGAGTTAAAAGCAGCCCT |
| sgNT1 | GCGAGGTATTCGGCTCCGCG |
| sgP53 Ex2 | CCTCGAGCTCCCTCTGAGCCAGG |
| sgPsen2-1 | TGCTCGCATTCATGGCCTCTC |
| sgPsen2-2 | GAGAGCACTGCCCAGTGGGT |

**Table S3:** **Genotyping primers used in the study**

| **Primer** | **Genotype** | **Sequence** |
| --- | --- | --- |
| oIMR0069 | C3(1)-TAg | GGACAAACCACAACTAGAATGCAGTG |
| oIMR0068 | C3(1)-TAg | CAGAGCAGAATTGTGGAGTGG |
| oIMR8745 | C3(1)-TAg Internal Positive Control | GTCAGTCGAGTGCACAGTTT |
| oIMR8744 | C3(1)-TAg Internal Positive Control | CAAATGTTGCTTGTCTGGTG |
